# Toward Rapid Sequenced-Based Detection and Characterization of Causative Agents of Bacteremia

**DOI:** 10.1101/162735

**Authors:** F. Curtis Hewitt, Stephanie L. Guertin, Krista L. Ternus, Kathleen Schulte, Dana R. Kadavy

## Abstract

Rapid pathogen diagnosis and characterization performed by metagenomic DNA sequencing may permit physicians to better target therapies in order to improve patient outcomes. To this end, a novel sample-to-answer workflow was assembled to enable rapid clinical detection of causative pathogens of bacteremia in whole blood utilizing metagenomic sequence data captured by the MinION. Rapid lysis, nucleic acid purification, host depletion, and genomic DNA library preparation permitted the detection of multiple bacterial and fungal agents spiked into whole blood, with sequencing commencing within 40 minutes of sample receipt. A hybrid detection strategy utilizing targeted PCR detection of specific pathogens of concern was adopted to improve overall sensitivity. As a proof of concept, primers for relatively long amplicons (~ 1800 bp) were selected to enable the specific detection of *Yersinia pestis*. The resulting amplicon library was spiked onto the same sequencing flow cell used to perform genomic sequencing, permitting simultaneous pathogen detection via both targeted and untargeted sequencing workflows. Sensitivities on the order of 1×10^6 cells/mL and 1x10^5 cells/mL were achieved for untargeted and targeted detection, respectively, of *Y. pestis* genomes spiked into whole blood. Bacterial and fungal species present in the ZymoBIOMICS Microbial Community Standard were also detected when spiked at similar levels. Variable quality of sequence reads was observed between the transposase-based and ligation-based library preparation methods, demonstrating that the more time consuming ligation-based approach may be more appropriate for the workflow described herein. Overall, this approach provides a foundation from which future point of care platforms could be developed to permit characterization of bacteremia within hours of admittance into a clinical environment.

**Author Summary:** Cases of bacteremia in the U.S. present a significant clinical challenge, especially due to rising rates of antimicrobial resistant strains. Rapid diagnosis of the etiologic pathogen and underlying drug resistance genetic signatures between the first and second antibiotic dose should improve patient outcomes and may permit physicians to better target antibiotic therapies without turning to broad spectrum antibiotics, which may further propagate resistant strains. The methods described herein have been developed to enhance the real time nature of the MinION sequencer. DNA sequencing and real time analysis begin within 40 minutes of sample receipt (as opposed to hours or days for common clinical nucleic acid extraction or blood culture techniques). The incorporation of sensitivity enhancements, such as methylation-based pulldown of human DNA or PCR targeted for pathogens of interest, ensures that this assay can detect bacterial blood infections at clinically relevant levels. The pathogen-agnostic aspect of the assay could one day allow clinicians to identify any unknown bacterial, fungal, or viral DNA in a sample. Ultimately, this study serves as an important step toward establishing a pipeline to rapidly detect and characterize pathogens present in whole blood.

## Introduction

High-throughput next generation sequencing platforms have not made a substantial impact on the human pathogen clinical diagnostics market despite the disruptive, transformative nature of the technology^1,2^. Beyond the challenge of achieving regulatory approval, the primary limitations associated with NGS for clinical diagnostics center largely on cost, throughput (especially time to answer), and the challenge of data analysis. As the cost of sequencing continues to decrease due to novel technologies and competition in the marketplace^3^, it becomes critical to increase assay speed and simplify data analysis in order to hasten the incorporation of NGS assays into the diagnostics marketplace.

The MinION™ (Oxford Nanopore Technologies, Oxford, UK; ONT) represents an ideal candidate to achieve rapid high throughput sequencing. The MinION performs single molecule sequencing via protein nanopores located within a sequencing flow cell that includes accompanying electronics^4^. This platform has significant appeal due to its simple library preparation, long read generation, flexible run times, and small footprint. Multiple studies have utilized the MinION to perform either long read shotgun sequencing capable of agnostic pathogen detection or targeted amplicon sequencing^5–8^. Sequencing accuracy and throughput have improved with the recent sequencing chemistries (e.g., R9.4), consistently generating > 80% single strand sequence identity and 5-10 GB of sequence data per 48 hour run at 450 bases/second, according to emerging reports^9^. While sequencing accuracy remains a potential drawback for clinical adoption, the rapid library prep (approximately 10 minutes) and real time nature of each sequencing run permit significant decreases in assay run time compared to other NGS platforms^7,8,10^. ONT also provides a suite of real time basecalling and metagenomics analysis tools, which promises to decrease the overall burden associated with bioinformatics analysis of NGS data^11^.

A quick sample preparation method is required to fully capitalize on the rapid library preparation and real time sequence analysis provided by the MinION. The workflow developed for this study incorporates several time saving modifications to minimize total sample preparation time. Analysis of whole blood presents a significant time savings compared to the generation of cell free fluids (i.e., blood plasma), albeit at the expense of generating significant host background sequence data. To ensure effective lysis of all potential pathogens present in a sample including viruses, gram positive and gram negative bacteria, fungal species, and hard-to-lyse bacterial and fungal spores, a mechanical method to shear cells (i.e., bead beating) was incorporated as a faster and more robust alternative to chemical lysis^10^. Further time savings were achieved via the use of expedited methods for DNA purification and pathogen enrichment. These magnetic bead-based pulldown methods represent a scalable method for purification when compared to column-based kits such as those produced by Qiagen^12^ or differential lysis kits for enrichment such as those produced by Molzym^13^.

To assess the assembled workflow, this study utilized a number of potential blood pathogens. Organisms utilized for testing included a naturally circulating biothreat pathogen (*Y. pestis*), various additional gram-negative bacteria (e.g., *Salmonella enterica*), several gram-positive bacteria (e.g., *Staphylococcus aureus*), as well as two fungal species. *Y. pestis,* which typically infects 1-20 individuals in the U.S. per year^14^, was selected as the main focus of this study due to the importance of strain-level identification which can be performed by the interrogation of genetic determinants including nucleotide variants and plasmid content. The diversity within the *Yersinia* genus also presents a challenge for metagenomic classification as accurate species identification must be performed to differentiate potential biothreat agents from less-pathogenic near neighbors (i.e., *Y. pseudotuberculosis, Y. enterocolitica*)^15,16^. The sample preparation methodology presented herein successfully identified the presence of a *Y. pestis* pathogen panel spiked into whole blood via both shotgun and targeted sequence analysis, albeit at relatively high titer levels. These results suggest a path forward for rapid sample preparation and metagenomic analysis at a throughput level (sample-to-answer in less than four hours) compatible with the needs of clinical diagnostics.

## Results and Discussion

### Workflow Design

The workflow was designed to achieve cell lysis, nucleic acid extraction, pathogen enrichment, and library preparation in as little time as possible. The underlying goal of this workflow time compression was to commence real time DNA sequencing and analysis as quickly as possible. The non-targeted workflow shown in Figure 1 allowed a prepared DNA library to be added to sequencing flow cells within 40 minutes of sample receipt (or approximately 20 minutes without pathogen enrichment). Significant time reductions compared to conventional whole blood DNA extraction methods were achieved via the utilization of the OmniLyse bead beater (ClaremontBio) for rapid cell lysis and magnetic bead-based approaches for DNA purification and enrichment. The rapid genomic transposase-based sequencing kit (SQK-RAD002; ONT) was used to prepare purified DNA for sequencing in approximately 10 minutes, and samples were then sequenced with the MinION. This portion of the workflow enables the collection of data from all DNA present in the sample, permitting detection of unexpected or novel pathogens. However, the sensitivity of this shotgun approach is limited by the amount of host background present in the sample.

**Fig 1.**
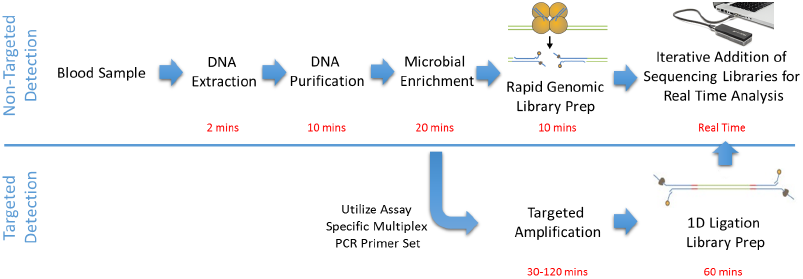
Workflow Overview. Production of a rapid workflow which includes modular enrichment steps (methylation-based enrichment, PCR). The workflow is compatible with small blood draw volumes (<250 μL). Real time analysis of shotgun sequencing data commences within approximately 40 minutes of sample receipt. Targeted detection commences within 2-4 hours dependent on the duration of target-specific PCR.

To improve sensitivity for specific pathogens, purified nucleic acids were also subject to targeted PCR and amplicon library preparation (Figure 1). The amplicon sequencing library was added directly to the same flow cell without washing away the genomic library, enabling parallel sequencing of the targeted and untargeted library. Real time basecalling (via cloud-based and local basecalling) and pathogen identification were performed using ONT-released tools. Third party metagenomic analysis tools were utilized to confirm results and provide additional confidence in pathogen classifications.

### Pathogen Enrichment for Agnostic Detection

Host background significantly confounds metagenomic analysis, especially in matrices such as whole blood, which contains > 10^6 nucleated cells/mL. Numerous host depletion or pathogen enrichment techniques have been reported for metagenomic sequencing of human matrices^12,17^. Here, we utilized the NEBNext^®^ Microbiome DNA Enrichment Kit (New England Biolabs; NEB), which utilizes magnetic beads to specifically bind eukaryotic methylation patterns (5-Methylcytosine in CpG dinucleotides). Enrichment was performed following nucleic acid purification, as shown in Figure 1. *Y. pestis* culture (Harbin 35 strain) was spiked into human whole blood at 2.58×10^6 cells/mL and processed via the pathogen-agnostic portion of the workflow (Figure 1) with or without enrichment. Sequencing of each library was carried out for approximately 16 hours with basecalling and metagenomic analysis via the ONT What’s In My Pot (WIMP) tool, which is based on Kraken^11^. A total of one read in the unenriched sample was attributed to *Yersinia* out of a total of 4,217 reads (Figure 2A). In the enriched sample, 57 reads were classified as *Yersinia* out of a total 8,649 reads, providing an overall 28-fold enrichment of *Yersinia* reads. The NEB enrichment process enabled detection of all bacterial species present in the Zymo mock community, which was spiked into human whole blood at 2.9×10^7 cells/mL (approximately 3×10^6 cells/mL of each species in the community) (Figure 2B). Fungal species present within the mock community were also observed following enrichment, as expected due to the lack of CpG methylation used to deplete host background DNA^18^. Despite low overall read counts across these experiments, potentially due to suboptimal DNA input or quality, these data suggest that host depletion plays a critical role in the overall sensitivity of this workflow.

**Fig 2.**
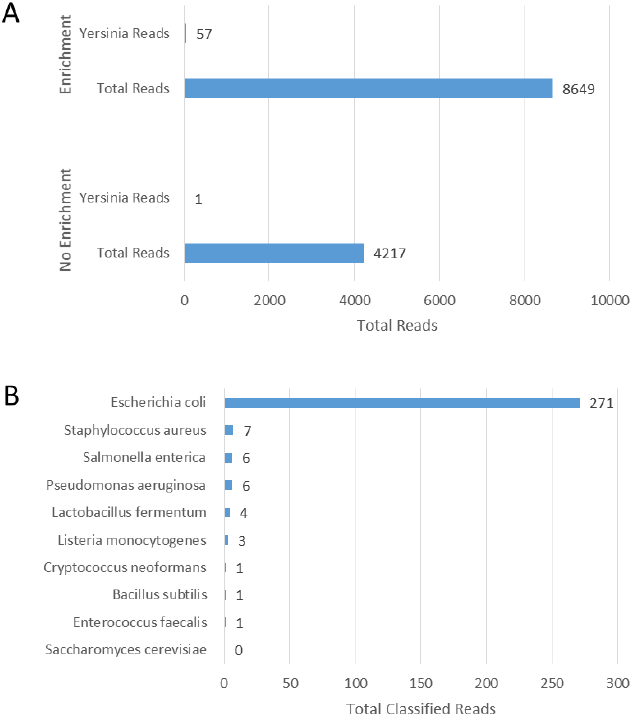
Detection of Pathogens in Whole Blood Requires Host Depletion. A) Metagenomic classification from shotgun sequence data of pathogens present in whole blood is significantly enhanced by depletion of host sequence. 0.66% of total reads are classified as Yersinia following sequence enrichment, compared to 0.023% of total reads when enrichment is not utilized corresponding to a 28-fold increase in the abundance of pathogen sequence. B) Representative detection of all species included in the Zymo mock community via shotgun sequencing and metagenomic classification following enrichment. Significant abundance of *E. coli* is likely due to laboratory contamination^33^.

### Pathogen Enrichment via Simultaneous PCR

MinION flow cells are flexible with regard to sequencing different libraries within each flow cell’s 48-hour sequencing lifetime. While loading multiple sequencing libraries from different samples/individuals is possible, carry-over between runs may create false positive detections that would be problematic for clinical diagnostics. However, preparing different libraries from the same sample overcomes the issue of contamination. As described in Figure 1, a hybrid shotgun/targeted approach was devised to permit targeted detection of high-consequence pathogens at greater sensitivities than possible with the untargeted pathway alone.

To test this method, a panel of primers specific to the *Y. pestis* genome and various plasmids was obtained (Table S1)^19–22^. The entire workflow described in Figure 1, including enrichment via the NEB kit, was performed as described above using human whole blood samples spiked with 2.58×10^6 cells/mL *Y. pestis* and approximately 3×10^6 cells/mL of each species in the Zymo mock community. A total of 1 μL (~ 20 ng) of the resulting DNA was subject to PCR using each primer set. PCR products were verified by gel electrophoresis and purified via AMPure XP beads (Beckman Coulter). Library preparation was carried out according to manufacturer’s instructions for ligation-based 1D sequencing. The ongoing genomic DNA sequence run was briefly stopped to permit loading of the amplicon library. Sequencing then resumed with data collection occurring at a significantly higher rate (> 1,000 reads per minute) than observed for the genomic library alone (Figure 3A). At this rate, significant coverage depth (greater than 30x coverage, sufficient for variant calling) of each amplicon was generated within minutes of sequencer re-initialization.

**Fig 3.**
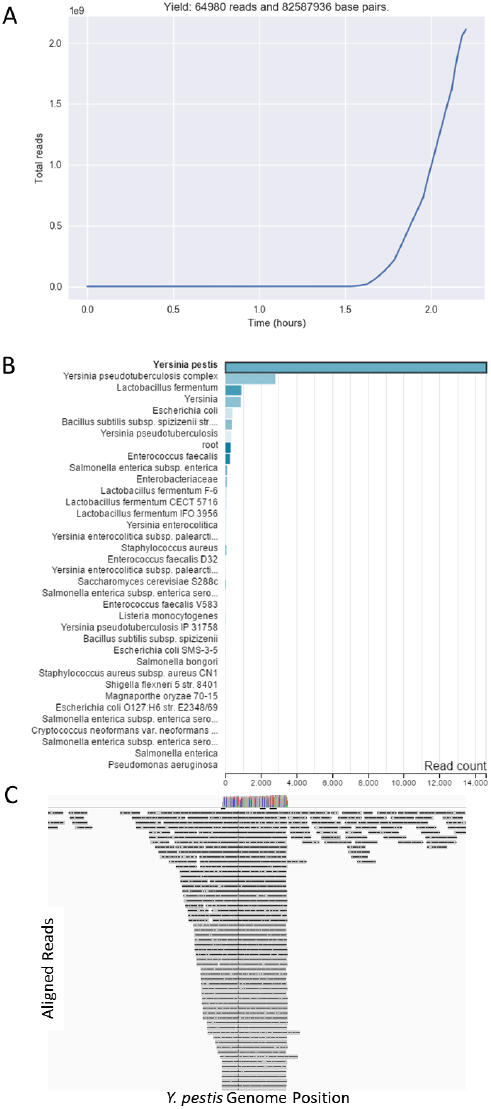
Hybrid Assay Provides Enhanced Pathogen-Specific Sensitivity without Compromising Detection of Background Pathogens. A) Total read count across duration of a hybrid workflow. Relatively few reads captured during the non-targeted portion of the assay (Time 0-1.6 hours) with significant increase in total read counts following the addition of the amplicon library. B) Metagenomic classification via WIMP following hybrid workflow. The majority of *Yersinia* reads are based on amplicon sequencing. Bacteria and fungi present in the Zymo mock community were identified as part of the genomic sequencing portion of the workflow. C) Representative visualization of reads aligned to the Harbin 35 reference genome. Significant depth of coverage of the amplicon sequence (> 1,000x coverage) observed at the expected location within the *Y. pestis* genome. Non-targeted sequences are also shown, interspersed across the genome at low coverage.

A total of 64,980 reads was successfully classified according to species, resulting from both the targeted and non-targeted aspects of the workflow. More than 14,000 reads, predominantly resulting from the amplicon sequencing portion of the workflow, were correctly attributed to *Y. pestis* (Figure 3B). The non-targeted portion of the workflow also detected *Yersinia* as well as every component of the Zymo mock community, albeit at very low sequence depth. Alignment of reads to the *Y. pestis* genome reveals the representative sequence depth along a portion of the reference genome compared to the significant sequence depth achieved via amplicon sequencing (Figure 3C).

### Variant Calling to Differentiate *Y. pestis* Strains

The genomic primer sets selected for testing encompass variants permitting the differentiation of *Y. pestis* strains. The amplicon targeted against the *ail* gene (Table S1) contains a single SNP which differentiates *Y. pestis* strain Harbin 35 from strain CO92^19^. The amplicon targeted against the *vasK* gene contains a 1 nt insertion in the Harbin 35 and CO92 strains, which can be used to differentiate these strains from strain KIM10^20^. To test whether these variants were correctly identified as part of the amplicon sequencing portion of the workflow, reads were aligned against the Harbin 35 genome and visualized using IGV (Figure 4). Surprisingly, the variant present near the *ail* gene (nt 2,998,385) did not conform to the reference genome. Of the 7,121 total reads at this site, 71% contained 2,998,385G, matching those reported in pathogenic strains such as CO92, and 26% of total reads contained the variant for the reference genome. These data suggest a potential mixture within the culture, as the 26% variant frequency is significantly higher than the sequencing error observed in the flanking region. The absence of the pgm locus was verified to ensure that this mixture did not include a pathogenic strain of *Y. pestis* (data not shown)^23^. Of the 3,141 total reads covering the *vasK* amplicon, 96% contained the variant in the reference genome (4,279,948C). Four hundred and sixteen reads contained a deletion at this location, as would be expected in the KIM10 strain. However, this is similar to the general indel rate in the flanking amplicon sequence, suggesting that this is an artifact of the current nanopore sequencing process. Overall, the current nanopore sequencing chemistry appears to support variant analysis for strain-level detection, albeit with error rates (especially indels^24^) which could lead to misclassifications.

**Fig 4.**
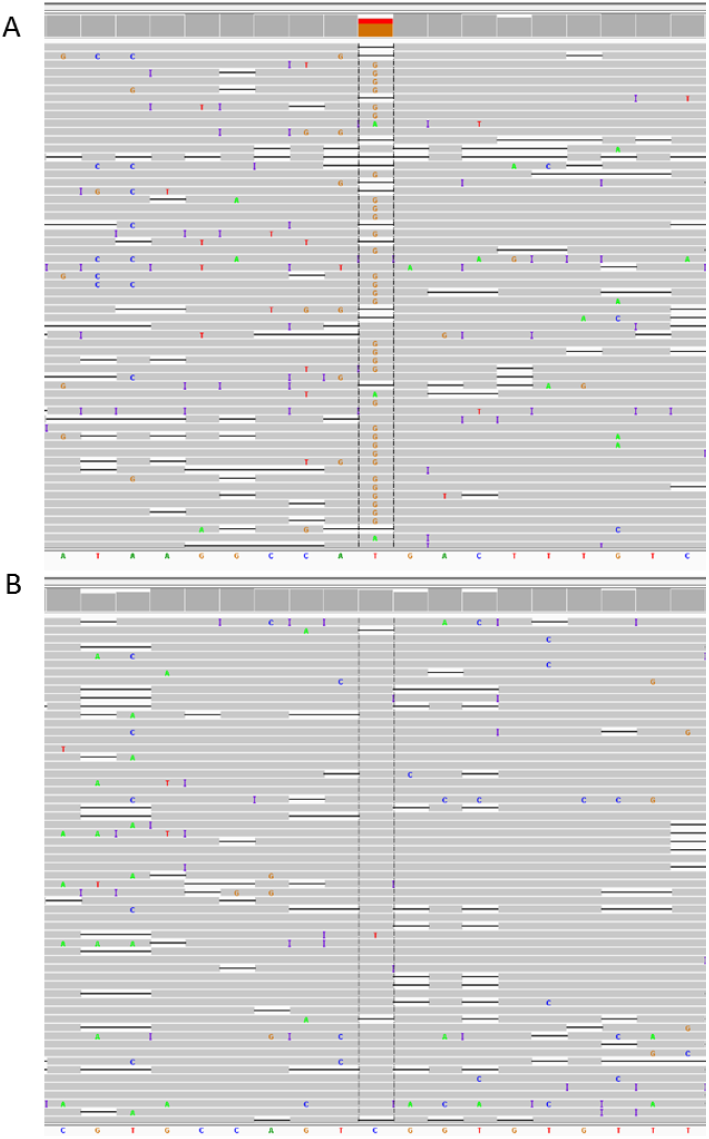
Visualizing Key Variants for Strain Identification. A) NC_017265.1:2,998,385. Mixture of T (26% of total reads, which corresponds to the reference genome) and G (71% of total reads, which corresponds to other *Y. pestis* strains including CO92). B) NC_017265.1:4,279,948. Majority of reads conform to the reference genome. Certain *Y. pestis* strains (e.g., Kim 10) contain a deletion of this base.

### Data Quality between Library Preparation Methods

Significant differences in quality score and sequence length distribution dependent on the library preparation method were consistently observed across four individual flow cells. Representative data is shown in Figure 5. In general, significantly higher quality scores were generated using 1D ligation-based library prep chemistry as opposed to the transposon-based rapid library prep method). This decrease in quality had a significant impact on the number of reads available for metagenomic analysis, especially for tools which impose a minimum quality score threshold prior to analysis, such as ONT’s WIMP tool. In line with the disparity in quality scores, the pass/fail ratio determined during basecalling was significantly higher for the ligation-based prep compared to the transposase chemistry.

**Fig 5.**
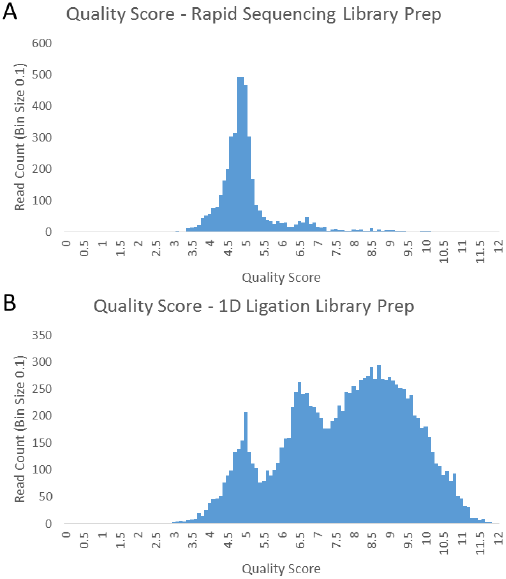
Relative Quality Scores between Transposase-and Ligation-Based Library Preparation Approaches. A) Quality scores resulting from the rapid genomic library prep sequencing kit. B) Quality scores resulting from the complete hybrid workflow.

It is possible that the utilization of input DNA quantities below manufacturer recommendations for rapid genomic sequencing contributed to lower quality scores; however, the observed pattern in sequencing quality was observed even when using highly purified pathogen DNA stocks at the recommended quantity. Under ideal conditions, the rapid library prep method yielded average and median QScores of 4.8 and 3.8, respectively. The ligation prep method yielded average and median QScores of 6.9 and 7.1, respectively. This data suggests that the shorter duration and simplified handling associated with the rapid library prep method may not sufficiently outweigh the decrease in sequence yield and quality observed as part of this workflow.

## Conclusion

The approach and results described herein demonstrates a potential pathway for rapid clinical metagenomic sequencing diagnostics while highlighting several key challenges. The ability to collect and analyze sequence data in real time holds disruptive potential for the clinical diagnostics marketplace. However, this potential can only be fully met if the sample and library preparation methods are similarly permissive to rapid, high confidence analysis. This study suggests that it is possible to perform metagenomic classification on whole blood samples without unnecessary complexity in upfront processing methods (e.g., plasma isolation). Further, sequencing of enriched samples can commence in as little as 40 minutes from sample receipt. Utilizing a parallel targeted analysis pathway via PCR still permits analysis to occur in less than four hours. Perhaps the most significant limitation in analysis speed remains the time required to transmit data to the cloud or physical storage for analysis – steps which could be separately enhanced by faster internet connections or greater local computational processing resources.

Based on the data presented here, achieving clinically relevant assay sensitivity appears to be a greater challenge than the accuracy of pathogen classification. While this study did not seek to perform a comprehensive limit of detection study on either the untargeted or targeted aspects of the workflow, it is clear that a significant amount of pathogen must be present in whole blood for detection to occur. Limits of detection for the workflow described herein likely fall within 1×10^5 to 1×10^6 cells/mL for metagenomic shotgun sequencing, with limits of detection for targeted sequencing likely one to two logs lower. While these figures are in line with peak titer numbers for many clinical pathogens causative of bacteremia, including *Y. pestis*^25^, this assay could lead to false negatives at lower titers or earlier stages of infection. These limits of detection also focus solely on pathogen identification and the presence/absence of related plasmids; in contrast, pathogen characterization (i.e., identifying drug resistance genotype variants) requires significantly higher sequencing coverage for high confidence genotyping. Additional input DNA or sequencing time would be required to achieve sufficient coverage for these types of characterization, unless the regions of interest within specific pathogens were incorporated into the targeted portion of the workflow.

Overall, this study presents an important initial step toward rapid detection of clinical pathogens in human whole blood. However, it is clear that significant gains in sensitivity must be achieved through increased sequencing throughput or pathogen enrichment. Anticipated gains in sequencing throughput communicated by ONT via chemistry improvements or parallel sequencing on larger platforms (e.g., GridION) should enhance sensitivity without substantially increasing sample-to-answer duration. Additional pathogen enrichment methods, such as plasma isolation to generate cell free fluids, or sequence capture, will also improve assay sensitivity^26,27^; However, these steps add additional time to sample processing and increase overall assay complexity, which may eventually limit the adoption of similar assays in laboratories which perform only CLIA-waived or moderately complex diagnostic tests. Software-based methods such as ONT’s Read Until are perhaps the most promising method to increase sensitivity without increasing sample-to-answer time or per-sample cost by selectively rejecting host background sequence reads during sequencing^28^. Continued development and optimization of this and similar assays hold the promise that rapid human clinical pathogen diagnostics via non-targeted sequence genotype assays are on the horizon.

## Materials and Methods

### Sample Preparation

Human whole blood (lithium heparin anticoagulant) was obtained from BioreclamationIVT. *Y. pestis* strain Harbin 35 (NR-639) was obtained through BEI Resources. ZymoBIOMICS™Microbial Community Standards were obtained from Zymo Research and consisted of the following organisms: *Pseudomonas aeruginosa, Escherichia coli, Salmonella enterica, Lactobacillus fermentum, Enterococcus faecalis, Staphylococcus aureus, Listeria monocytogenes, Bacillus subtilis, Saccharomyces cerevisiae, Cryptococcus neoformans*. The concentrations of all cultures were determined via hemocytometer across three replicates. The *Y. pestis* stock had a concentration of 1.29×10^8 cells/mL and the Zymo mock community stock had a total concentration of 2.42×10^9 total cells/mL (equivalent to approximately 3×10^8 cells/mL of each bacterial species). To spike blood samples, the indicated concentration of bacterial culture was added to 500 μL whole blood and vortexed gently for ~ 10 seconds.

### Lysis and Nucleic Acid Purification

Sample lysis was performed with the OmniLyse (ClaremontBio) per manufacturer’s instructions. The OmniLyse cartridge was pre-rinsed with approximately 500 μL of molecular biology grade water (ThermoFisher) for one minute. Lysis of 250 μL of human whole blood spiked with bacteria was performed for two minutes. Initial nucleic acid purification was performed via AMPure XP beads (BeckmanCoulter). An additional 1.8 volumes of beads were added to the lysed solution. After binding for 10 minutes at ambient temperature, the pellet was washed 3x with 500 μL fresh 70% ethanol. DNA was eluted in 40 μL of molecular biology grade water. Purity and DNA concentration were measured via Nanodrop (ThermoFisher). Pathogen enrichment was carried out using the NEBNext^®^ Microbiome DNA Enrichment Kit (New England Biolabs). MBD2-Fc beads were pre-incubated with Protein A beads prior to each experiment to reduce total assay time. Remaining enrichment steps followed manufacturer’s protocol with final elution in 40 μL of molecular biology grade water.

### Amplification

Purified nucleic acid was subject to PCR using Phusion DNA polymerase (New England Biolabs). One μL of template DNA was added to each 50 μL PCR reaction. Individual primer sets are listed in Table S1. PCR products were purified using AMPure XP beads as per manufacturer instructions and eluted in 40 μL molecular biology grade water. Concentrations were measured via Nanodrop. Products were later verified by gel electrophoresis (1% agarose in 1X TAE buffer). The PCR products were mixed to produce a cocktail with 1 µg of total DNA (0.2 ng of each of five PCR products) for library preparation.

### MinION library preparation and sequencing

MinION library preparations were prepared according to the appropriate protocols. Genomic sequencing was performed using the Rapid Sequencing of genomic DNA kit (SQK-RAD002). Amplicon sequencing was performed using the 1D Amplicon Sequencing kit (SQK-LSK108). The maximum amount of input DNA volume (7.5 μL) was added for rapid genomic sequencing; 1 μg of total DNA was utilized as input for amplicon sequencing library preparation. Prepared libraries were loaded onto Spot-On flow cells (SpotON Flow Cell Mk I – R9.4). Amplicon libraries were added to the same flow cell 90-120 minutes after addition of the genomic library. The sequencer was reinitialized and run for an additional 6-24 hours depending on the sequencing run. Basecalling was performed using the local version of the albacore version 1.1.2 basecaller.

### Data analysis

Real time analysis was performed using the What’s In My Pot (WIMP) tool provided by Oxford Nanopore Technology’s Metrichor^11^. Basecalled reads were sampled based on sequencing time for comparative analysis between experiments as indicated. Reads binned according to pass and fail were combined prior to additional analysis. Poretools^29^ version 0.6.0 was used to generate fastq files for the MinION 1D reads. As WIMP does not contain a human reference in the database, One Codex)^5^ was used as a comparator and to properly identify host/background sequence. Sequence alignments were generated using Graphmap (v0.22)^30^. Resulting SAM alignment files were converted to BAM format using SAMtools (v1.4)^31^. Visual alignment of reads and confirmation of variant calls was performed using IGV version 2.3^32^. Reads were aligned against the *Yersinia pestis* biovar Medievalis strain Harbin 35 genome (accession number CP001608.1).

## Acknowledgements

The authors would like to thank Oxford Nanopore Technologies for providing reagents for this work. The authors would also like to thank Dr. Charles Chiu for his technical input as part of this effort and the University of Texas Medical Branch for their recommendations on *Y. pestis* primers. The following reagent was obtained through BEI Resources, NIAID, NIH: *Yersinia pestis*, Strain Harbin 35, NR-639.

## Author Contributions

Conceived and designed the experiments: FCH KLT DRK. Performed the experiments: FCH KS KLT. Analyzed the data: SLG FCH KLT. Wrote the paper: FCH SLG.

## Data Availability

Raw DNA sequence reads have been deposited in the NCBI Sequence Read Archive under accession numbers SAMN07286086 and SAMN07286087.

